# Hydroxyl-radical scavenging activity of hydrogen does not significantly contribute to its biological function

**DOI:** 10.1101/2021.03.13.435216

**Authors:** Qinjian Li, Fei Xie, Yang Yi, Pengxiang Zhao, Xin Zhang, Xiaokang Zhang, Xujuan Zhang, Xuemei Ma

**Author notes:** Co-first author (Qinjian Li, Fei Xie). Corresponding author: Xuemei Ma.

## Abstract

Since Ohsawa et al. reported a biological antioxidant function of hydrogen in 2007, researchers have now shown it to exert protective effects in a wide range of human and animal disease models. Clinical observations and scientific arguments suggest that a selective scavenging property of H_2_ cannot adequately explain the beneficial effects of hydrogen. However, there is no experiment challenging the original published data, which suggested that molecular hydrogen dissolved in solution reacts with hydroxyl radicals in cell-free systems. Here we report that a hydrogen-saturated solution (0.6 mM) did not significantly reduce hydroxyl radicals in the Fenton system using 1 mM H_2_O_2_. We replicated the same condition as Ohsawa’s study (i.e. 5 μM H_2_O_2_), and observed a decrease in ^•^OH radicals in both the H_2_-rich and N_2_-rich solutions, which may be caused by a decreased dissolved oxygen concentration. Finally, we determined the effect of hydrogen on a high-valence iron enzyme, horseradish peroxidase (HRP), and found that hydrogen could directly increase HRP activity in a dose-dependent manner. Overall, these results indicate that although H_2_ and ^•^OH can react, the reaction rate is too low to have physiological function. The target of hydrogen is more complex, and its interaction with enzymes or other macro-molecules deserve more attention and in-depth study.

## Introduction

In 2007, Ohsawa et al. [1] reported neuroprotective effects of molecular hydrogen in a rat model of cerebral infarction. They further demonstrated that H_2_ could act as a therapeutic antioxidant by selectively reducing hydroxyl radicals without disturbing important physiological reactive oxygen species (ROS). Data generated since have demonstrated potential benefits of H_2_ in over 170 different human and animal disease models [2]. For example, H_2_ has beneficial effects in both type 1 [3] and type 2 diabetes, insulin resistance [4,5], experimental liver injury [6], acute oxidative stress in focal brain ischemia/reperfusion injury [1], acute and chronic stress [7], and organophosphorus pesticide-induced neurotoxicity [8]. H_2_ was also shown to have some anti-cancer effects [9,10], and alleviate nephrotoxicity induced by anti-tumor drugs [11]. Several clinical trials, such as in patients with potential metabolic syndrome [12,13], Parkinson’s disease [14], rheumatoid arthritis [15], mild cognitive impairments [16], and others, as reviewed previously [2,17], further demonstrate hydrogen’s potential as a medical gas.

These therapeutic effects are associated with beneficial changes in several transcription factors (e.g. Nf-κβ, Nrf2, NFAT, etc.) and subsequent regulation of anti-oxidant and anti-inflammatory mediators (e.g. glutathione, superoxide dismutase, cytokines, TNF-α, etc.) [17]. However, the underlying mechanism(s) and primary target(s) remain elusive, as many of these molecules are simply passenger molecules that are changed secondarily by modulation of their upstream regulators [2].

To date, there are only four reported primary actions of molecular hydrogen that may help explain some, but not all of its ubiquitous biological effects. The first, and most widely cited, is direct ^•^OH radical scavenging by H_2_ (i.e. H_2_ + ^•^OH → H_2_O) [1]. However, this hypothesis does not adequately explain the many effects of H_2_. In fact, a companion paper published in the same issue of the *2007-Nature Medicine* has questioned this hypothesis theoretically [18], indicating that there was no conclusion on this issue at that time. Although more than 1000 original articles have been published in the field of molecular hydrogen from 2007, and the specific hydroxyl radical scavenging effect of hydrogen has been repeatedly proposed in these articles, only a few articles have questioned this hypothesis again [19], and no conclusive experimental evidence has been presented to confirm or refute this hypothesis so far. A second hypothesis, showing the oxidation of deuterium gas *in vitro*, suggests an iron species-dihydrogen interaction with the iron sulfur clusters within mitochondrial complex I [20]. A third in vitro experiment, suggests an interaction between hydrogen and certain enzymes, such as increasing the activity of acetylcholine esterase (AChE) [8]. The fourth hypothesis is of an *in vitro* experiment where it was demonstrated that H_2_ gas could suppress the auto-oxidation of linoleic acid, and subsequently modify lipid mediators thus affecting Ca^2+^ signaling and gene expression [21]. However, these proposed mechanisms require further investigation to falsify and/or reproduce their results, and to determine how much, and to what extent they can address the ubiquitous therapeutic effects of only small amounts of molecular hydrogen used *in vivo*.

In this study, we directly investigated the first hypothesis to determine if H_2_ significantly reacts with ^•^OH radicals or if it could be explained by other study confounds (e.g. decreased dissolved O_2_). We also investigated another potential target of molecular hydrogen, and report the preliminary positive *in vitro* results, which may further elucidate the mechanisms by which H_2_ exerts its biological benefits.

## Materials and Methods

### Reagents

Hydrogen peroxide (30%), ferrous (II) perchlorate, K_3_Fe(CN)_6_, were purchased from Sigma, 2-[6-(4’-hydroxy) phenoxy-3H-xanthen-3-on-9-yl] benzoate (HPF) from Molecular Probes^®^ and Horse radish peroxidase (HRP) was purchased from Aladdin^®^. Solutions were prepared with deionized water, which was purified using a Milli-Q water purification system for laboratory tests.

### Hydroxyl radical producing methods

The composition of ^•^OH -radical generating system is shown in Table 1. H_2_ or N_2_ gas was bubbled in ultrapure water beyond the saturated level under 0.4 MPa of pure hydrogen pressure for 30 min, and the resultant H_2_ or N_2_-rich water was then used to prepare ^•^OH-producing system under atmospheric pressure. We determined the H_2_ concentration with a hydrogen electrode (Unisense A/S, Aarhus, Denmark) in each experiment. The concentration of dissolved oxygen in H_2_-rich water (3.65 mg/L), N_2_-rich water (3.50 mg/L) and normal water without treatment (Air) (8.75 mg/L) was determined using an oxygen electrode (Thermo Fisher Scientific, USA).

**Table 1.** The composition of the ^•^OH-generation reaction system

| Reaction System (50 $\mu\text{L}$) | Composition with $\text{H}_2$ (0.8 mM) or $\text{N}_2$-rich water | |
| --- | --- | --- |
| Fenton reaction | $\text{H}_2\text{O}_2$<br>(5 $\mu\text{M}$ or 1 mM) | Ferrous (II) perchlorate<br>(0.1 mM) |

### Detection of ^•^OH formation by hydroxyphenyl fluorescein

To detect the reaction of H_2_ and ^•^OH, the Fenton reaction system as described above was used in the presence of HPF (0.4 μM). The fluorescence intensity of HPF was measured at 535 nm with excitation at 485 nm by using a Victor 1420 Multilabel Counter (Wallac, Perkin-Elmer, Wellesley, MA, USA).

### Horseradish peroxidase (HRP) enzyme activity test

We incubated solutions containing 0.4 μM HPF, 10 μM H_2_O_2_ with H_2_ or N_2_-rich water at different concentrations (85%, 50% and 25%). The reaction was initiated by adding 0.2 μg/mL HRP. The fluorescence of HPF was measured at 535 nm with excitation at 485 nm by using a Victor 1420 Multilabel Counter (Wallac, Perkin-Elmer, Wellesley, MA, USA).

### Reduction of Fe(III) by H_2_

The fluorescent intensity of HPF was also used to test the possible reduction of ferric iron (Fe^3+^) by H_2_, FeCl_3_ (0.1 mM), H_2_O_2_ (5 μM), HPF (0.4 μM) and either H_2_ or N_2_-rich water were mixed together. The fluorescent intensity of HPF was measured at 535 nm with excitation at 485 nm by using a Victor 1420 Multilabel Counter (Wallac, Perkin-Elmer, Wellesley, MA, USA). For K_3_Fe(CN)_6_ reaction, 2 mM K_3_Fe(CN)_6_, 0.4 μM HPF and H_2_ or N_2_ rich water was mixed, followed by HPF fluorescence measurement.

### Statistical analysis

Results are expressed as means ± S.E.M. for each experiment. For single comparisons, we used an unpaired two-tailed Student’s *t* test; for multiple comparisons, we used ANOVA. p < 0.05 was considered to be statistically significant.

## Results

### The effect of hydrogen on the ^•^OH production in cell-free *in vitro* system

In Ohsawa’s study, they produced ^•^OH radicals by the Fenton reaction and semi-quantified the levels of ^•^OH by HPF fluorescence [1]. A previous study showed that both the presence of oxygen and the [Fe^2+^]/H_2_O_2_ ratio may have an impact on ^•^OH production in the Fenton reaction [22]. To determine whether the H_2_ induced ^•^OH decrease is caused by the reduced oxygen concentration, N_2_ was used to induce similar anaerobic conditions and the ^•^OH formation was detected. To further determine whether a different [Fe^2+^]/H_2_O_2_ ratio could affect ^•^OH production in the Fenton reaction, we used a H_2_O_2_ concentration of either 5 μM (the same as Ohsawa’s study) or 1 mM. These concentrations represent a high (20:1) and low (1:10) [Fe^2+^]/H_2_O_2_ ratio, respectively. The ^•^OH formation in Fenton reaction was assessed via HPF fluorescence detection. As expected, both H_2_ and N_2_ decreased the HPF fluorescent signal of ^•^OH radicals in the presence of 5 μM H_2_O_2_ (Fig. 1A and B). However, in the presence of 1 mM H_2_O_2_, there was neither a decrease nor an increase in ^•^OH formation in either the H_2_ or N_2_ group (Fig. 1C and D).

**Fig. 1.**
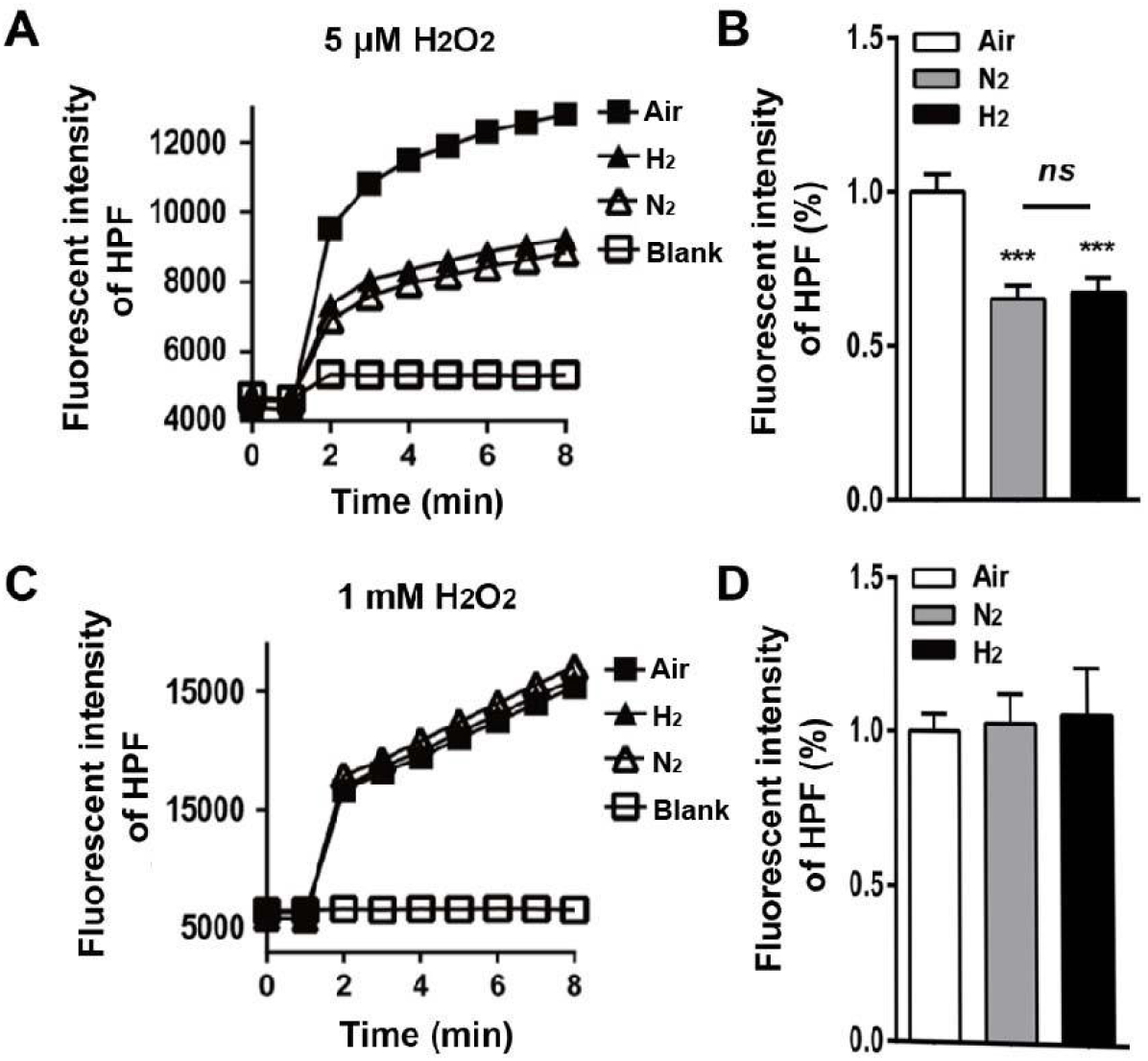
The effect of hydrogen on ^•^OH formation in Fenton reaction assessed by HPF fluorescence. (A) and (C) are the time course of ^•^OH formation in the presence of 5 μM (A) or 1 mM H_2_O_2_ (C). (B) and (D) are the quantification of the HPF fluorescence intensity (5 μM H_2_O_2_ (B), n = 8; 1 mM H_2_O_2_ (D), n = 10). ***p < 0.001, *ns* p > 0.05.

### Effect of hydrogen on horseradish peroxidase activity

We tested the activity of HRP, based on the HPF fluorescence intensity, in the presence of 10 μM H_2_O_2_ in water with or without H_2_ or N_2_. The results showed that both hydrogen and nitrogen treatment greatly increased HPF signals in a dose-dependent manner (Fig. 2A), and the HPF signals in each dilution in H_2_ group were markedly higher than that in the N_2_ group (Fig. 2B). These results indicated that hydrogen and nitrogen treatment may enhance the HRP activity, an affect which may not be attributed to the removal of oxygen.

**Fig. 2.**
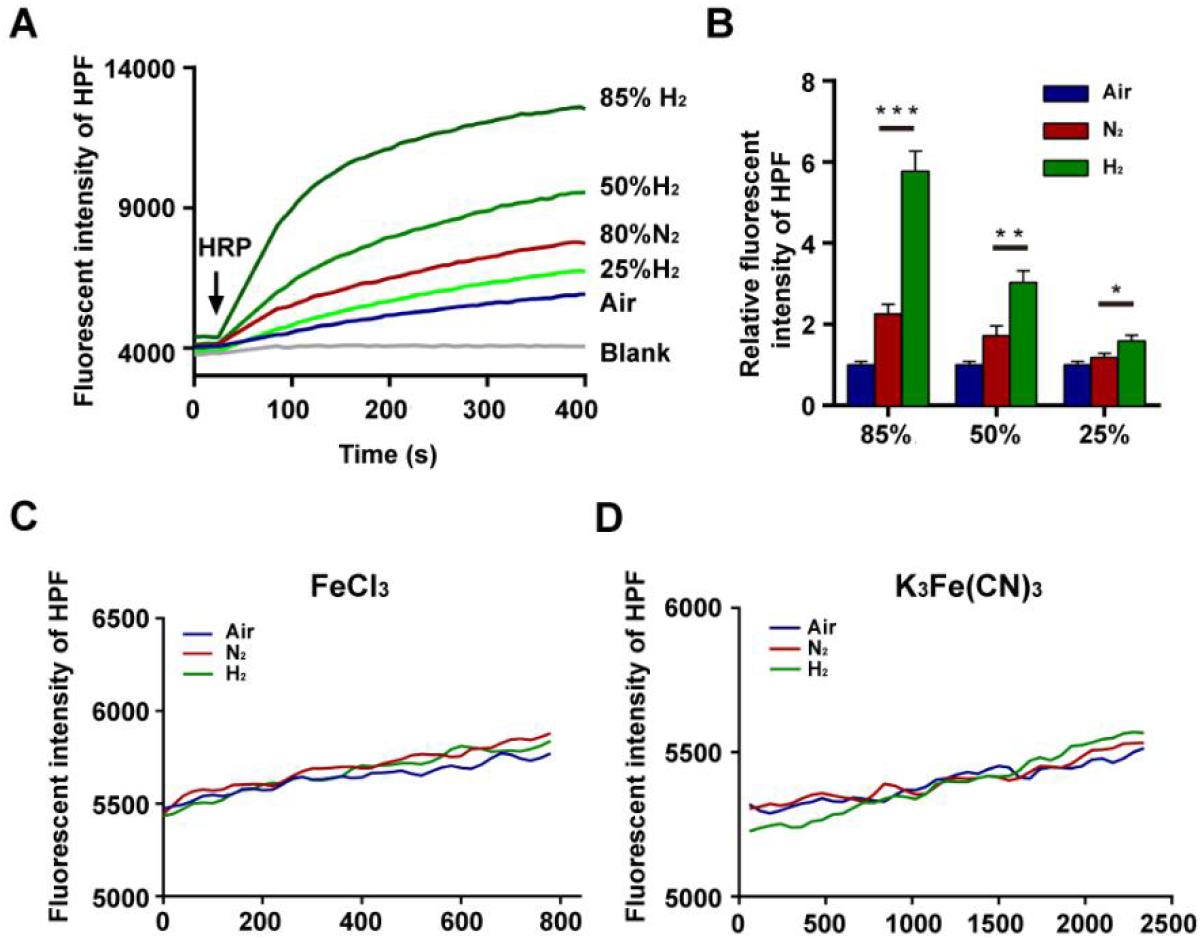
The possible effects of hydrogen on Fe(III) and on HRP activity were determined by HPF florescence. (A) is the representative time course of HPF fluorescence at different concentration of H_2_ and 85%N_2_. Baseline showed HPF fluorescence in absence of HRP. (B) is the quantification of HPF fluorescence at 300 seconds after adding HRP. (C) and (D) is the representative time course of HPF fluorescence in the presence of FeCl_3_ and K_3_Fe(CN)_6_, respectively. *p<0.05, **p<0.01, ***p<0.001.

### Effect of hydrogen on the redox state of Ferric iron (Fe^3+^)

To further confirm whether H_2_ could possibly reduce Fe(III), the HPF fluorescence based method was used. The results showed no observable reduction in either FeCl_3_ (Fig. 2C) or K_3_Fe(CN)_6_ (Fig. 2D).

## Discussion

Since Ohsawa et al. published the first paper in *Nature Medicine*, the effects of hydrogen have been extensively reported using more than 170 different human and animal diseases models [2]. The idea that H_2_ acts as a therapeutic antioxidant via directly interacting with ^•^OH radicals was first proposed in Ohsawa’s study [1]. They evaluated the ^•^OH scavenging activity of H_2_ in an *in vitro* cell-free Fenton reaction by HPF fluorescence based method, and found that H_2_ reduced the HPF fluorescent signal of ^•^OH radicals in a dose-dependent manner. One of the widely interpretations of this study, is that hydrogen can selectively scavenge ^•^OH, which account of its therapeutic and antioxidant-like effects. This perspective has also been questioned [18,19], due to two main concerns. The first is the cellular concentration of H_2_ only reaches the micromolar range, but in order to be an effective biological ^•^OH scavenger, the required concentration should be in the millimolar range, and in some organs (e.g. brain) the H_2_ level may never even increase after drinking H_2_-rich water [23]. The second is that the observed rate constant for the H_2_ and ^•^OH reaction (i.e. 4.2×10^7^ M^-1^ s^-1^), is likely too low to effectively compete with the other reactions between ^•^OH and numerous cellular targets such as proteins or lipids, whose rate constants are on the order of 10^9^ or 10^10^ M^-1^ s^-1^ [19].

In the present study, to determine whether H_2_ significantly reacts with ^•^OH radicals, we evaluated the ^•^OH scavenging activity of H_2_ in an *in vitro* cell-free Fenton reaction by using HPF fluorescence based method. The fluorescent intensity of HPF was used as the indicator of the levels of ^•^OH radicals. In the presence of 1 mM H_2_O_2_ concentration, there was no reduction of ^•^OH radicals in neither the H_2_ nor the N_2_ group. We note that in Ohsawa’s study, they used a very low H_2_O_2_ concentration (5 μM), which is close to the *in vivo* physiological level, but started the reaction with 0.1 mM Fe^2+^. This leads to an excess of Fe^2+^ in conditions with 5 μM H_2_O_2_ which is oxidized by the oxygen in the air-saturated media but not in the H_2_ purged solutions where oxygen is removed or diminished. In our study, the same concentration of H_2_O_2_ in the Fenton reaction was also used. In contrast to the higher level of H_2_O_2_, the results showed that the levels of ^•^OH radicals were significantly decreased in H_2_ group compared to the control, which was consistent with Ohsawa’s study. However, this marked reduction in ^•^OH radical formation was also observed in the N_2_ group. It was reported that besides changes in O_2_ concentration, the [Fe^2+^]/H_2_O_2_ ratio also impacts ^•^OH production in the Fenton reaction [22]. When the [Fe^2+^]/H_2_O_2_ ratio is high (> 1.5), the ^•^OH production in anaerobic conditions is markedly lower than in aerobic conditions [22]. In our study, in the presence of 5 μM H_2_O_2_, the [Fe^2+^]/H_2_O_2_ ratio reached to 20:1. We speculate that in the high [Fe^2+^]/H_2_O_2_ ratio condition, the decrease in ^•^OH formation may be attributed to the reduced oxygen concentration. A previous study also showed that in a low [Fe^2+^]/H_2_O_2_ ratio condition (< 1), the decrease in oxygen concentration had no significant effect on the ^•^OH production [22]. This may explain why in the presence of 1 mM H_2_O_2_ ([Fe^2+^]/H_2_O_2_ ratio: 1:10), there was neither a decrease nor an increase in ^•^OH formation in either the H_2_ or the N_2_ group. Our results demonstrate that under these conditions with infeasible supraphysiological concentrations of H_2_O_2_ (1 mM) and H_2_ (≈ 0.6 mM) there was no reduction in ^•^OH radicals in the H_2_ group. Thus, we propose that although H_2_ can react with ^•^OH radicals at an observed rate constant of 4.2×10^7^ M^-1^ s^-1^ [19], hydroxyl radical scavenging of hydrogen does not significantly contribute to its biological function.

Next, we hypothesized that, because direct ^•^OH-radical scavenging activity of H_2_ likely does not have biological significance, H_2_ must interact with other biomolecules. Besides ^•^OH radicals, HPF can also be oxidized by various enzymes containing high-valence iron, such as horseradish peroxidase (HRP) [24]. HRP, a related heme enzyme of cytochrome c peroxidase and cytochrome P450, has an analogous high-valence iron intermediate, but does not release hydroxyl radicals [25]. To determine whether hydrogen could directly modulate the activity of an enzyme containing high-valence iron, we tested the activity of HRP based on the HPF fluorescence intensity with or without H_2_ or N_2_ in a cell-free *in vitro* system. Our study clearly showed that hydrogen treatment greatly increased HPF signals in a dose-dependent manner, which cannot be fully explained by removal of oxygen, as is evidenced by significantly higher level of HPF signals in the H_2_ group compared to the N_2_ group. These results suggest that high-valence iron containing enzyme(s) could be one of the potential targets of hydrogen.

Our previous study was the first to report an interaction between molecular hydrogen and biological macromolecules [8]. We found that molecular hydrogen could directly increase the activity of acetylcholinesterase (AChE). In combination with the evidence provided in the present study, we propose that the biological macromolecules, especially the enzymes, might be a potential target of hydrogen involved in its biological functions.

Lastly, a previous study has also investigated the notion that H_2_ may exert its beneficial effects by reducing Fe(III) centers, which are oxidized during oxidative stress. However, they reported that neither hemes nor iron–sulfur clusters were reduced in cytochrome P450cam, myoglobin, and putidaredoxin [19]. Liu *et al.* showed that in addition to hydroxyl radicals, HPF can also be oxidized by the high-valence iron and K_3_Fe(CN)_6_ [24]. In this study, we determined the possible reductive effect of hydrogen on K_3_Fe(CN)_6_ and FeCl_3_ by measuring the intensity of HPF fluorescence. Consistently, no significant effect was observed both in the presence of either H_2_ or N_2_, which indicated that molecular hydrogen does not reduce Fe(III) centers.

In conclusion, in this study we provided evidence that the selective-scavenging activity of hydrogen for ^•^OH radicals may not be the principal factor that contributes to its biological functions. Instead, other mechanisms/targets such as biological macromolecules, especially enzymes, might be responsible for its therapeutic effects.

## Conflict of interest statement

We declare that we do not have any commercial or associative interest that represents a conflict of interest in connection with the work submitted.

## Acknowledgments

We want to gratefully acknowledge the helpful comments and suggestions from Professor Shigeo Ohta, Tyler W. LeBaron and Yingxian Li. This research was supported by National Natural Science Foundation of China (grant number: 31500828) and Basic Research Foundation of Beijing University of Technology (grant number: 015000514314005).

## Notes

### Competing Interest Statement

The authors have declared no competing interest.

### Summary of Updates

The order of the legend of Figure1A and 1C ("N2", "H2") has been revised as "H2", "N2".

## Reference

[1] I. Ohsawa, M. Ishikawa, K. Takahashi, M. Watanabe, K. Nishimaki, K. Yamagata, K. Katsura, Y. Katayama, S. Asoh, S. Ohta, Hydrogen acts as a therapeutic antioxidant by selectively reducing cytotoxic oxygen radicals, Nat. Med. 13 (2007) 688–694.

[2] M. Ichihara, S. Sobue, M. Ito, M. Hirayama, K. Ohno, Beneficial biological effects and the underlying mechanisms of molecular hydrogen – comprehensive review of 321 original articles, Med. Gas Res. 5 (2015) 12.

[3] H. Amitani, A. Asakawa, K. Cheng, M. Amitani, K. Kaimoto, M. Nakano, M. Ushikai, Y. Li, M. Tsai, J. Li, M. Terashi, H. Chaolu, R. Kamimura, A. Inui, Hydrogen Improves Glycemic Control in Type1 Diabetic Animal Model by Promoting Glucose Uptake into Skeletal Muscle, Plos One 8 (2013) e53913.

[4] X. Zhang, J. Liu, K. Jin, H. Xu, C. Wang, Z. Zhang, M. Kong, Z. Zhang, Q. Wang, F. Wang, Subcutaneous injection of hydrogen gas is a novel effective treatment for type 2 diabetes, J. Diabetes Investig. 9 (2018) 83–90.

[5] S. Kajiyama, G. Hasegawa, M. Asano, H. Hosoda, M. Fukui, N. Nakamura, J. Kitawaki, S. Imai, K. Nakano, M. Ohta, T. Adachi, H. Obayashi, T. Yoshikawa, Supplementation of hydrogen-rich water improves lipid and glucose metabolism in patients with type 2 diabetes or impaired glucose tolerance, Nutr. Res. 28 (2008) 137–143.

[6] H. Sun, L. Chen, W. Zhou, L. Hu, L. Li, Q. Tu, Y. Chang, Q. Liu, X. Sun, M. Wu, H. Wang, The protective role of hydrogen-rich saline in experimental liver injury in mice, J. Hepatol. 54 (2011) 471–480.

[7] Q. Gao, H. Song, X. Wang, Y. Liang, Y. Xi, Y. Gao, Q. Guo, T. LeBaron, Y. Luo, S. Li, X. Yin, H. Shi, Y. Ma, Molecular hydrogen increases resilience to stress in mice, Sci. Rep. 7 (2017) 9625.

[8] T. Wang, L. Zhao, M. Liu, F. Xie, X. Ma, P. Zhao, Y. Liu, J. Li, M. Wang, Z. Yang, Y. Zhang, Oral intake of hydrogen-rich water ameliorated chlorpyrifos-induced neurotoxicity in rats, Toxicol. Appl. Pharm. 280 (2014) 169–176.

[9] J. Runtuwene, H. Amitani, M. Amitani, A. Asakawa, K. Cheng, A. Inui, Hydrogen-water enhances 5-fluorouracil-induced inhibition of colon cancer, PeerJ. 3 (2015) e859.

[10] M. Dole, F. Wilson, W. Fife, Hyperbaric hydrogen therapy: a possible treatment for cancer, Science 190 (1975) 152–154.

[11] N. Nakashima-Kamimura, T. Mori, I. Ohsawa, S. Asoh, S. Ohta, Molecular hydrogen alleviates nephrotoxicity induced by an anti-cancer drug cisplatin without compromising anti-tumor activity in mice, Cancer Chemoth. Pharm. 64 (2009) 753–761.

[12] G. Song, M. Li, H. Sang, L. Zhang, X. Li, S. Yao, Y. Yu, C. Zong, Y. Xue, S. Qin, Hydrogen-rich water decreases serum LDL-cholesterol levels and improves HDL function in patients with potential metabolic syndrome, J. Lipid Res. 54 (2013) 1884–1893.

[13] A. Nakao, Y. Toyoda, P. Sharma, M. Evans, N. Guthrie, Effectiveness of hydrogen rich water on antioxidant status of subjects with potential metabolic syndrome-an open label pilot study, J. Clin. Biochem. Nutr. 46 (2010) 140–149.

[14] A. Yoritaka, M. Takanashi, M. Hirayama, T. Nakahara, S. Ohta, Pilot study of H2 therapy in Parkinson’s disease: a randomized double-blind placebo-controlled trial, Mov. Disord. 28 (2013) 836–839.

[15] T. Ishibashi, B. Sato, M. Rikitake, T. Seo, R. Kurokawa, Y. Hara, Y. Naritomi, H. Hara, T. Nagao, Consumption of water containing a high concentration of molecular hydrogen reduces oxidative stress and disease activity in patients with rheumatoid arthritis: an open-label pilot study, Med. Gas Res. 2 (2012) 27.

[16] K. Nishimaki, T. Asada, I. Ohsawa, E. Nakajima, C. Ikejima, T. Yokota, N. Kamimura, S. Ohta, Effects of molecular hydrogen assessed by an animal model and a randomized clinical study on mild cognitive impairment. Curr. Alzheimer Res. 14 (2017).

[17] G. Nicolson, G. Mattos, R. Settineri, C. Costa, R. Ellithorpe, Clinical effects of hydrogen administration: from animal and human diseases to exercise medicine. Int. J. Clin. Med. 7 (2016) 32–76.

[18] K.C. Wood, M.T. Gladwin, The hydrogen highway to reperfusion therapy, Nature Med. 13 (2007) 673–674.

[19] J. Penders, R. Kissner, W.H. Koppenol, ONOOH does not react with H2: Potential beneficial effects of H2, as an antioxidant by selective reaction with hydroxyl radicals and peroxynitrite, Free Radical Bio. Med. 75 (2014) 191–194.

[20] R. Hyspler, A. Ticha, H. Schierbeek, A. Galkin, Z. Zadak, The evaluation and quantitation of dihydrogen metabolism using deuterium isotope in rats, PLoS One 10 (2015) e0130687.

[21] K. Iuchi, A. Imoto, N. Kamimura, K. Nishimaki, H. Ichimiya, T. Yokota, S. Ohta, Molecular hydrogen regulates gene expression by modifying the free radical chain reaction-dependent generation of oxidized phospholipid mediators. Sci. Rep. 6 (2016) 18971.

[22] S.Y. Qian, G.R. Buettner, Iron and dioxygen chemistry is an important route to initiation of biological free radical oxidations: an electron paramagnetic resonance spin trapping study, Free Radical Bio. Med. 26 (1999) 1447–1456.

[23] C. Liu, R. Kurokawa, M. Fujino, S. Hirano, B. Sato, X. Li, Estimation of the hydrogen concentration in rat tissue using an airtight tube following the administration of hydrogen via various routes. Sci. Rep. 4 (2014) 5485.

[24] Y. Liu, J.A. Imlay, Cell death from antibiotics without the involvement of reactive oxygen species, Science 339 (2013) 1210–1213.

[25] T.L. Poulos, Heme Enzyme Structure and Function, Chem. Rev. 114 (2014) 3919–3962.

